# Direct stimulation of anterior insula and ventromedial prefrontal cortex disrupts economic choices

**DOI:** 10.1101/2023.12.07.570630

**Authors:** Romane Cecchi, Antoine Collomb-Clerc, Inès Rachidi, Lorella Minotti, Philippe Kahane, Mathias Pessiglione, Julien Bastin

**Affiliations:** Univ. Grenoble Alpes, Inserm, U1216, CHU Grenoble Alpes, Grenoble Institut Neurosciences, 38000 Grenoble, France; Neurology Department, University Hospital of Grenoble, Grenoble, France; Motivation, Brain and Behavior (MBB) team, Paris Brain Institute, Pitié-Salpêtrière Hospital, Paris, France; Université de Paris, Paris, France

## Abstract

Neural activities within the ventromedial prefrontal cortex and anterior insula are associated with economic choices. However, whether these brain regions are causally related to these processes remains unclear. To address this issue, we leveraged rare intracerebral electrical stimulation (iES) data in epileptic patients. We show that opposite effects of iES on choice depend on the location of stimulation on a dorso-ventral axis within each area, thus demonstrating dissociable neural circuits causally involved in accepting versus avoiding challenges.

## Main text

Take a risk or play it safe? Although it is widely accepted that risky decisions are influenced by the desirability of potential outcomes, the neural mechanisms underlying such choices remain poorly understood. Functional imaging and intracranial electrophysiological evidence suggest that the ventromedial prefrontal cortex (vmPFC) and the anterior insular cortex (aIns) play critical roles in shaping these decisions. Accordingly, increased pre-stimulus neural activity in the vmPFC and aIns has been shown to promote and temper risk-taking, respectively, by overweighting the prospects of monetary gain or loss (Cecchi et al., 2022; Vinckier et al., 2018). Although the timing of these activities suggests a possible causal relationship between these brain regions and choice behavior, the correlational nature of the techniques used to date prevents conclusive establishment of this causal relationship.

In parallel, another line of research has shown that neural activity in both the vmPFC (De Martino et al., 2013; Lebreton et al., 2015; Lopez-Persem et al., 2020; Shapiro & Grafton, 2020) and aIns (Hebart et al., 2016; Molenberghs et al., 2016; Pereira et al., 2020; Vaccaro & Fleming, 2018) is related to participants’ confidence in their own decisions. Furthermore, confidence judgments are in turn influenced by monetary prospects (Lebreton et al., 2018, 2019), thus suggesting that the neural codes for value and confidence may not be independent.

To date, causal studies investigating the involvement of the vmPFC and aIns in risky decision-making and confidence judgments have provided limited evidence due to several limitations. First, the deep anatomical location of these brain regions has posed challenges in targeting these areas using non-invasive stimulation methods (Howard et al., 2020; Manuel et al., 2019; Spagnolo et al., 2019). Additionally, lesion studies have yielded inconsistent results on risk-taking behavior that may be attributed to variations in the scope and precise location of brain lesions (Bechara et al., 1994, 1999; Clark et al., 2008; Leland & Grafman, 2005; Reber et al., 2017; Sanfey et al., 2003; Shiv et al., 2005; Weller et al., 2007, 2009). Finally, intracranial electrical stimulation (iES) studies have rarely examined these regions (Duong et al., 2023; Yih et al., 2019) and have not combined iES with cognitive paradigms designed to tease apart precise decision-making processes.

To address these challenges, we conducted an experiment using intracranial electrical stimulation (iES) over human cortex to determine whether iES can disrupt choice behavior and also confidence judgments using a previously validated accept/reject task (Cecchi et al., 2022; Vinckier et al., 2018). Previous research has indicated that iES of the ventral aIns elicits disgust behavior or depressed affect (Caruana et al., 2011; Jezzini et al., 2012; Krolak-Salmon et al., 2003; Singh et al., 2021), whereas iES of the dorsal aIns elicits overwhelming ecstatic sensations or ingestive behavior, such as chewing and swallowing (Bartolomei et al., 2019; Jezzini et al., 2012; Nencha et al., 2022; Picard et al., 2013). Expanding on this evidence, we posit that harnessing the high spatial resolution of the iES will allow us to uncover functional circuits causally related to choice behavior.

The effect of intracranial electrical stimulation (iES) on choice and confidence was assessed in a cohort of 15 participants (aged 34.9 ± 2.7 years, including 7 females, **Table S1**) with epilepsy who had intracerebral electrodes implanted to localize their epileptogenic zone. During the choice phase of the task, participants were instructed to decide whether to accept the upcoming challenge for the stakes offered or to decline the offer and play for minimal monetary gains/losses. Additionally, participants were required to provide prospective confidence judgments regarding their task performance (**Figure 1a-b**).

**Figure 1.**
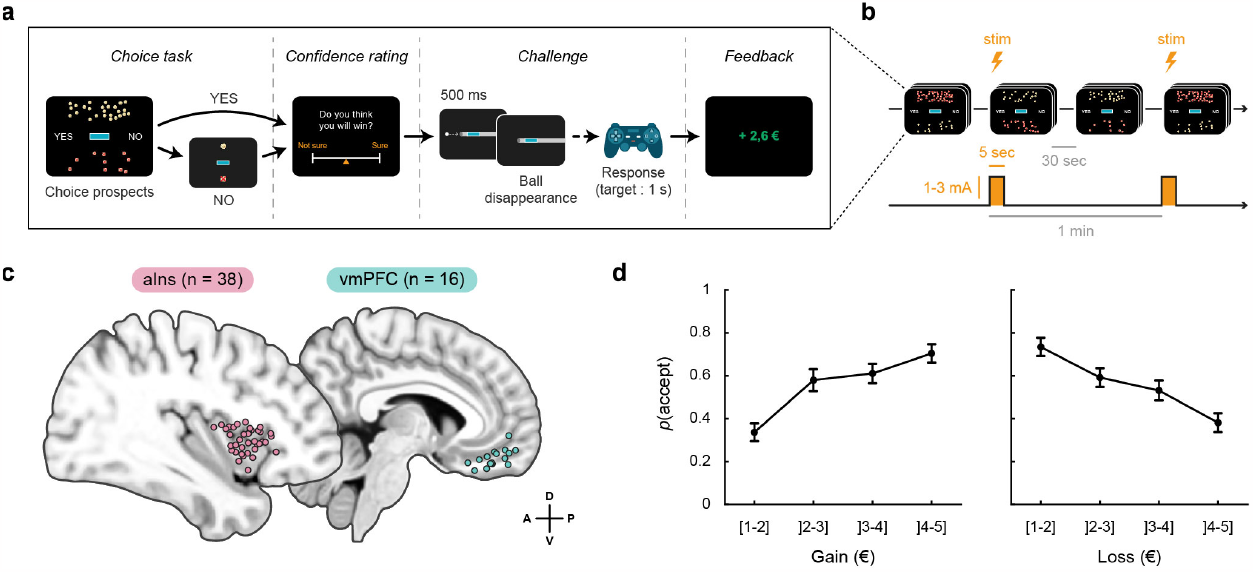
Experimental design and choice behavior. **(a)** Trial structure. Each trial consisted of a choice task and a confidence rating, followed by a challenge and finally feedback. In the choice task, subjects had to decide whether to accept or reject a given offer by considering gain prospects (represented by a bunch of regular 10-cent coins) and loss prospects (crossed out 10-cent coins). The challenge consisted of stopping a moving ball, which was invisible when entering the gray tunnel, within the blue target (difficulty level) in the center of the screen. **(b)** Overview of the stimulation procedure. In each session, 14 trials with intracranial electrical stimulation (iES) alternated with 14 trials without iES. The nature of the first trial (with or without iES) was randomized so that subjects remained blind to experimental conditions. A 30-second interval was observed between each trial onset, resulting in a 1-minute interval between each stimulation. During iES trials, stimulation was delivered for 5 seconds as pulses of 500 ms width at a frequency of 50 Hz and an amplitude of 1, 2, or 3 mA. **(c)** Anatomical location of the anterior insula (aIns; purple) and ventromedial prefrontal cortex (vmPFC; green) stimulation sites retained for analysis on the standard Montreal Neurological Institute (MNI) template brain. Anterior (A), posterior (P), dorsal (D) and ventral (V) directions are indicated. Note that stimulation sites have been aggregated in the mediolateral direction (x-axis) for visualization purposes. See **Supplementary Figure 1** for a more detailed location of each site. **(d)** Choice behavior in trials without iES. Acceptance probability is plotted as a function of the monetary prospects (gain and loss). Circles are binned data averaged across stimulation sites. Error bars represent inter-subject S.E.M.

We examined the effect of iES across a maximum number of intracerebral sites, resulting in data collected from 54 stimulation sites in which iES was applied to either the aIns (**Figure 1c**; n = 38 stimulation sites from 13 participants) or to the vmPFC (**Figure 1c**; n = 16 stimulation sites from 9 participants). Behavioral results from the trials without iES were consistent with previous observations in this task (**Figure 1d**). Specifically, participants’ acceptance rate was positively correlated with the amount of potential monetary gain and negatively correlated with the potential monetary loss (β_gain_ = 0.11 ± 0.03, t_772_ = 4.24, p = 3.10^−5^; β_loss_ = −0.09 ± 0.02, t_772_ = −4.26, p = 2.10^−5^; logistic mixed-effects regression). In contrast, the challenge difficulty did not significantly modulate choice behavior (β_diff_ = 7.10^−3^ ± 0.02, t_772_ = 0.33, p = 0.741), as we experimentally set it to levels where participants’ performance was close to chance. Furthermore, confidence ratings reported during trials without iES were positively associated with the probability of accepting the challenge (β_conf_ = 0.39 ± 0.17, t_758_ = 2.23, p = 0.026; logistic mixed-effects regression), indicating reliable metacognitive judgments.

Preliminary analysis of behavioral data suggested that iES might differentially affect behavior depending on the anatomical location of iES application within the aIns or vmPFC brain regions (**Figure 2a**). To quantify this, we examined whether the spatial coordinates of iES could predict choice (in the Montreal Neurological Institute space). The results confirmed that iES location had a significant impact on the choices made in both brain regions.

**Figure 2.**
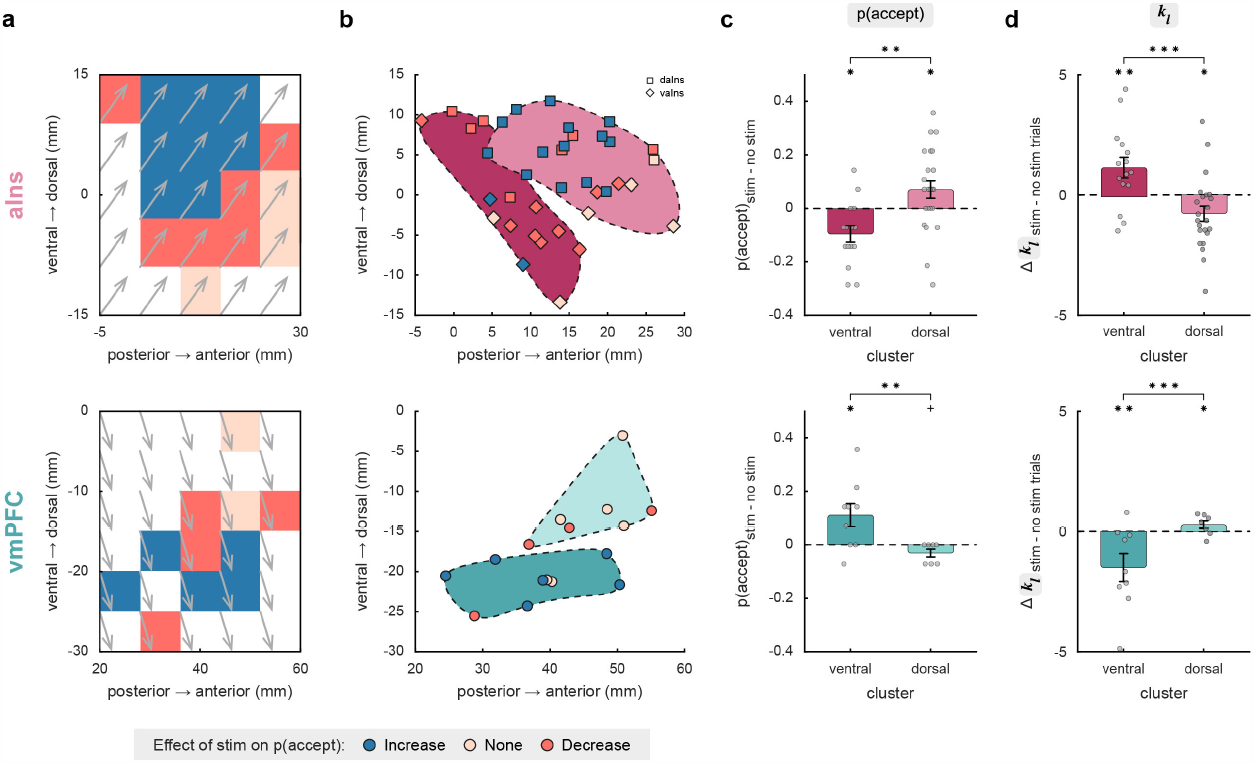
Stimulation effects on acceptance probability and computational mechanisms. **(a)** Spatial gradient of the effect of intracranial electrical stimulation (iES) on acceptance probability along the anteroposterior (y-axis) and ventrodorsal (z-axis) directions in the anterior insula (aIns; top) and ventromedial prefrontal cortex (vmPFC; bottom). Gray arrows indicate the direction of the gradient. **(b)** Clusters obtained by k-means clustering are plotted along the anteroposterior and ventrodorsal axes. **(c)** Effects of iES on clusters. (d) Effects of iES on participants’ sensitivity to potential losses (k_l_). In panels (c) and (d), dots represent the individual differences in acceptance probability and model weight (i.e., posterior parameter) k_l_ between trials with and without iES for each stimulation site. Bars and error bars represent mean and SEM, respectively, across stimulation sites. *p < 0.05, **p < 0.01, ***p < 0.001, + indicates a trend with p = 0.056. Stars between bars represent the interaction between stimulation condition and anatomical cluster obtained using a linear mixed-effects model with subjects and stimulation sites as random effects. Stars above a single bar represent post-hoc analyses.

In the aIns, there was a significant effect along the antero-posterior (*y*) axis (β_Y_ = 0.04 ± 0.02, t_34_ = 2.15, p = 0.039) and a trend along the ventro-dorsal (*z*) axis (β_Z_ = 0.04 ± 0.02, t_34_ = 1.86, p = 0.072), whereas in the vmPFC, the ventro-dorsal (*z*) localization of iES had a significant influence on choices made (β_Z_ = -0.10 ± 0.03, t_12_ = -3.71, p = 3.10^−3^). These findings were replicated using K-means clustering to the data (**Figure 2b**). This confirmed the existence of two distinct areas within each brain region (aIns: a postero-ventral cluster with n = 15 sites vs. an antero-dorsal cluster with n = 23 sites; vmPFC: a ventral cluster with n = 9 sites vs. a dorsal cluster with n = 7 sites).

In the following, we further specify how iES impacted economic choices (**Figure 2c**). In the aIns, we observed a significant interaction between stimulation condition and anatomical clusters (*F*_1,72_ = 11.6, p = 1.10^−3^; linear mixed-effects model). Post-hoc analyses revealed that iES decreased risk-taking when applied to the ventral part of the aIns (*F*_1,72_ = 6.7, p = 0.012), whereas iES increased risk-taking when applied to the dorsal part of the aIns (*F*_1,72_ = 5.4, p = 0.023). Similarly, we found a significant interaction between stimulation condition and vmPFC clusters (*F*_1,28_ = 10.1, p = 4.10^−3^). In contrast to the aIns results, post-hoc analyses revealed an opposing pattern, where iES increased the probability of accepting the challenge when applied to the ventral part of the vmPFC (*F*_1,28_ = 6.5, p = 0.016), while it tended to decrease risk-taking when applied to the dorsal part of the vmPFC (*F*_1,28_ = 4.0, p = 0.056).

Next, we investigate the computational mechanisms through which iES disrupted choice behavior by fitting the choice data with a model based on expected utility theory (Cecchi et al., 2022; Vinckier et al., 2018). Our findings revealed that iES modulated participants’ sensitivity to potential losses (captured by the parameter *kl*) in both brain regions (**Figure 2d**), but not their sensitivity to potential gains (**Supplementary Figure 2**). Specifically, we observed a significant interaction between stimulation condition and clusters regarding the effect on *kl* in both the aIns (*F*_1,72_ = 15.3, p = 2.10^−4^) and vmPFC (*F*_1,28_ = 13.6, p = 9.10^−4^). Subsequent post-hoc analyses revealed that iES applied to the ventral part of the aIns increased *kl* (*F*_1,72_ = 7.6, p = 8.10^−3^) while iES applied to its dorsal part decreased *kl* (*F*_1,72_ = 6.3, p = 0.014). Again, an opposite dorso-ventral gradient was identified within the vmPFC, whereby iES in its ventral part decreased *kl* (*F*_1,28_ = 8.9, p = 6.10^−3^), while iES in its dorsal part increased participants’ sensitivity to monetary loss prospects (*F*_1,28_ = 5.2, p = 0.030).

To exclude the possibility that the previous results were partly biased by a double-dipping issue (since K-means clusters were identified to delineate functional subregions within either aIns or vmPFC), we next defined dorsal or ventral functional subregions within the aIns or vmPFC using individual anatomical landmarks. For the anterior insula, we used Destrieux atlas (Destrieux et al., 2010) to distinguish between iES sites located within the short insular gyri and anterior circular sulcus of the insula (corresponding to the ventral aIns) and sites located within the superior circular sulcus (corresponding to the dorsal aIns). iES applied to anatomically defined subregions of the aIns differentially affected participants’ choices in the same directions as in the previous analysis (**Supplementary Figure 3;** interaction: *F*_1,72_ = 12.4, p = 8.10^−4^). Specifically, iES increased risky decisions in the dorsal aIns (post-hoc: *F*_1,72_ = 6.6, p = 0.012), whereas iES decreased risk-taking in the ventral aIns (post-hoc: *F*_1,72_ = 6.6, p = 0.012). Similar results were found for participants’ sensitivity to loss (interaction: *F*_1,72_ = 11.9, p = 1.10^−3^), where iES in the dorsal aIns decreased *kl* (*F*_1,72_ = 5.3, p = 0.025), while iES in the ventral aIns increased *kl* (*F*_1,72_ = 6.5, p = 0.013). Unfortunately, the spatial sampling from the anatomically-defined dorsal part of the vmPFC (namely the suprarostral sulcus; SU-ROS) was not sufficient (n = 3) to perform a similar analysis for the vmPFC. Nevertheless, upon evaluating the effect of iES solely on the ventral part of the vmPFC (**Supplementary Figure 4**; recordings within the superior rostral sulcus; ROS-S), we observed that iES increased risky-choices (*F*_1,20_ = 4.5, p = 0.046; linear mixed-effects model) and decreased participants’ sensitivity to potential losses (*kl*; *F*_1,20_ = 5.0, p = 0.037), consistent with results based on statistically defined subregions (K-means clustering approach).

Finally, we examined the influence of iES on confidence levels. In the aIns, there was a significant negative main effect of iES on confidence scores (**Figure 3**; *F*_1,70_ = 9.9, p = 2.10^−3^; linear mixed-effects model), with no dissociable effect of iES on confidence in its dorsal and ventral parts (**Supplementary Figure 5**; non-significant interaction effect *F*_1,70_ = 3.10^−3^, p = 0.96). This means that applying iES over the entire aIns significantly reduced participants’ confidence compared to trials without iES. This effect was not explained by a possible impact of iES on performing the challenge (*F*_1,72_ = 0.21, p = 0.65). In the vmPFC, no significant interaction (*F*_1,28_ = 1.0, p = 0.33) or main effect of stimulation (*F*_1,28_ = 0.04, p = 0.83) and cluster (*F*_1,28_ = 2.10^−13^, p = 1) was found, suggesting that iES on this brain region did not affect participants’ confidence levels (**Figure 3**).

**Figure 3.**
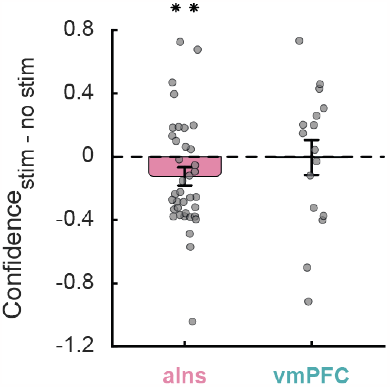
Stimulation effects on confidence. Dots represent individual difference in confidence ratings between trials with and without intracranial electrical stimulation for each stimulation site in the anterior insula (aIns; left) and ventromedial prefrontal cortex (vmPFC; right). Bars and error bars respectively represent mean and SEM across stimulation sites. **p < 0.01. The stars represent the main effect of stimulation, obtained when examining the effect of stimulation conditions and clusters on confidence ratings using a linear mixed-effects model with subjects and stimulation sites as random effects.

## Discussion

An emerging consensus is that the vmPFC and aIns may play opposing roles during economic decisions, mood, or learning, with neural activity in the vmPFC associated with increased risk-seeking, pleasantness, reward-based learning and good mood (Engelmann & Tamir, 2009; Gueguen et al., 2021; Tobler et al., 2007; Venkatraman et al., 2009; Xue et al., 2009), whereas neural activity in the aIns has been linked to risk-averse decisions, unpleasantness, punishment-avoidance, and bad mood (Cecchi et al., 2022; Kuhnen & Knutson, 2005; Paulus et al., 2003; Pessiglione et al., 2006; Rolls et al., 2008; Rudorf et al., 2012; Venkatraman et al., 2009; Vinckier et al., 2018). Our findings demonstrate the causal role of aIns and vmPFC functional subregions during choices and confidence. Whereas the dorsal vmPFC and ventral aIns reduced risk-taking by increasing participants’ sensitivity to monetary loss prospects, the ventral part of vmpFC and dorsal aIns increased risk-taking by decreasing participant’s loss sensitivity. Conversely, iES applied to the aIns decreased confidence, whereas iES did not disrupt confidence judgments in the vmPFC.

In the aIns, we have demonstrated that applying iES to the anterior portion of the superior circular sulcus led to an underestimation of loss prospects, resulting in an increase in risk-taking behavior, while stimulation of the ventral aIns (short insular gyri and anterior circular sulcus) led to an overestimation of loss prospects, resulting in a decrease in risk-taking behavior. This results extends to humans previous findings from mice showing distinct areas within the insula controlling approach vs. avoidance behavior (Peng et al., 2015; Wang et al., 2018). Our findings also bridges with stimulation studies in non-human primates where stimulation of the ventral aIns reduced approach behaviors in appetitive contexts (Saga et al., 2019) and was associated with negative behaviors such as disgust (Caruana et al., 2011; Jezzini et al., 2012; Krolak-Salmon et al., 2003) or depression (Singh et al., 2021). Conversely, iES on the dorsal aIns has been more commonly associated with positive behaviors such as ingestion in monkeys (Jezzini et al., 2012) or even ecstatic feelings in rare case studies in humans (Bartolomei et al., 2019; Nencha et al., 2022; Picard et al., 2013). Our findings bridge the gap between previous human and preclinical evidences by demonstrating within a single cohort of participants dissociable effects of iES within aIns by combining iES with an economic choice paradigm. Interestingly, iES on aIns decreased confidence (independently from iES localization), thus suggesting anatomo-functional dissociation of choice and confidence in this region. This is to our knowledge the first causal evidence for a critical role of aIns during metacognitive judgments, thus extending previous findings in which neural activity of aIns was correlated with confidence (Hebart et al., 2016; Molenberghs et al., 2016; Pereira et al., 2020; Vaccaro & Fleming, 2018).

In the vmPFC, our results indicate that stimulation of the superior rostral sulcus (ROS-S) led to an underestimation of loss prospects, resulting in an increase in risk-taking. Since existing studies investigating the effects of stimulation of this area have often reported null results (Fox et al., 2018, 2020; Selimbeyoglu & Parvizi, 2010), we propose that combining iES with cognitive paradigms designed to tease apart precise decision-making functions paves the way for future research aiming at better specifying the functions of the vmPFC. Although our findings suggest the existence of subterritories within the vmPFC, we had a limited statistical power to investigate the effects of iES on the (anatomically defined) dorsal part of the vmPFC (i.e., the suprarostral sulcus, SU-ROS), such that future work is needed to further specify the precise boundary between the dorsal vs. ventral portions of the vmPFC. Finally, despite the existing evidence associating the vmPFC with confidence judgments across various tasks (De Martino et al., 2013; Gherman & Philiastides, 2018; Hoven et al., 2022; Lebreton et al., 2015; Lopez-Persem et al., 2020; Morales et al., 2018), we failed to demonstrate a causal role of the vmPFC during confidence. Although negative findings should be interpreted with caution, we can only speculate that metacognitive judgments may involve a higher level of variability or noise, which, together with the fact that they were also temporally delayed to the iES onset compared to choices, may have impeded the emergence of a statistically significant effect of iES on the vmPFC regarding confidence.

Finally, it is important to acknowledge limitations inherent to this study. iES data relied on individuals with severe drug-resistant epilepsy, raising questions about the generalizability of our findings to the broader population, while the clinical context also limited the number of trials with iES for obvious ethical considerations. Yet, the advantage of iES was its ability to target specific brain regions with high spatial precision, allowing us to differentiate between ventral and dorsal parts of aIns or vmPFC. Achieving such sub-regional specificity would be challenging with other causal techniques available in human cognitive neuroscience.

To conclude, our study represents a step forward in elucidating the causal contribution of opponent circuits involving the vmPFC and aIns, revealing the existence of dorso-ventral subregions during value-based decisions. Given the involvement of these circuits in several neuropsychiatric disorders, our findings offer valuable insights which will be useful for the emerging development of closed-loop brain stimulation approaches to better alleviate symptoms that are otherwise resistant to existing treatments.

## Supplementary material

**Supplementary Figure 1:**
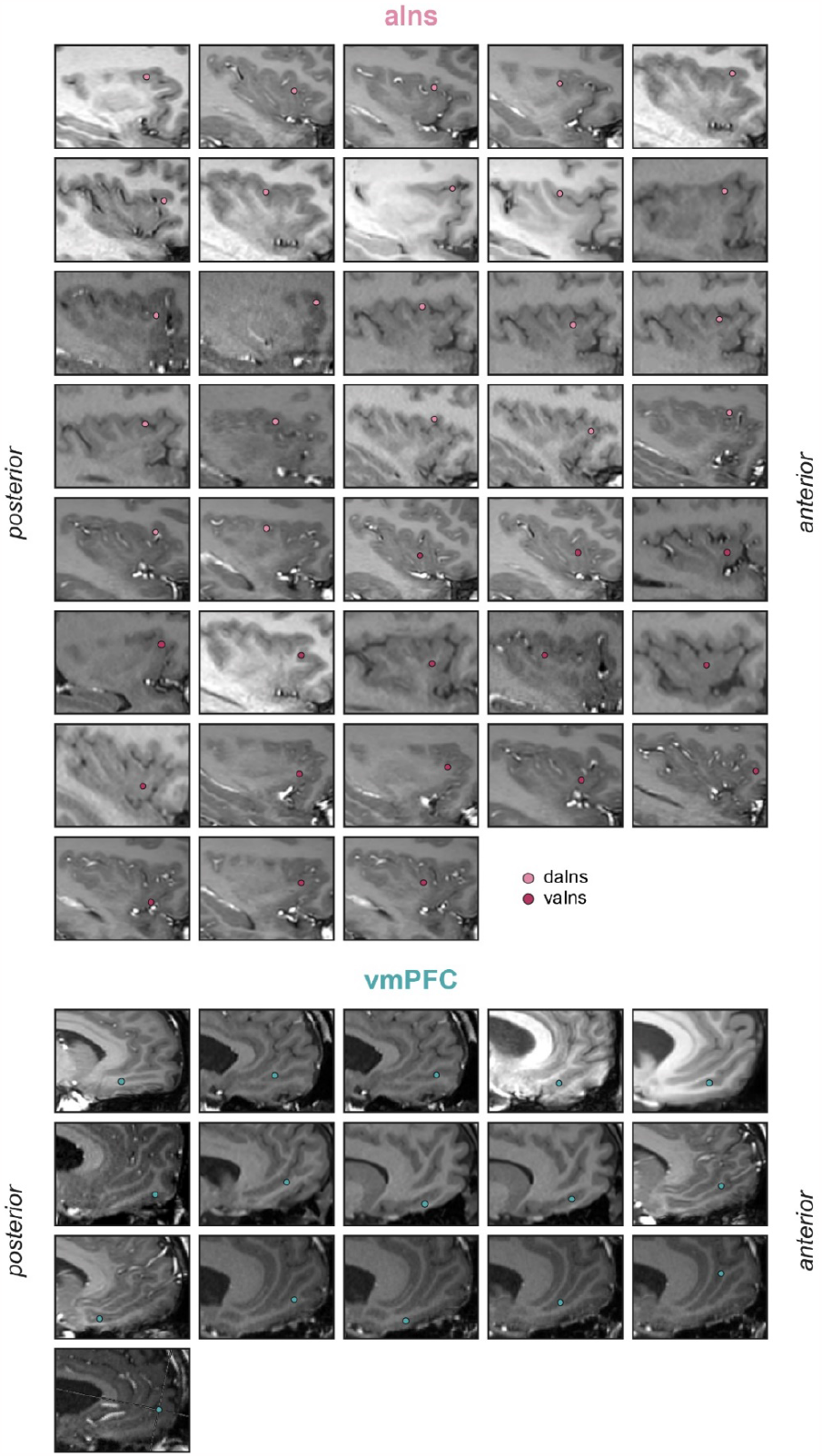
Location of stimulation sites on individual MRI brain slices. The locations of each stimulation site in the dorsal anterior insula (daIns), ventral anterior insula (vaIns), and ventromedial prefrontal cortex (vmPFC) are displayed on sagittal slices.

**Supplementary Figure 2:**
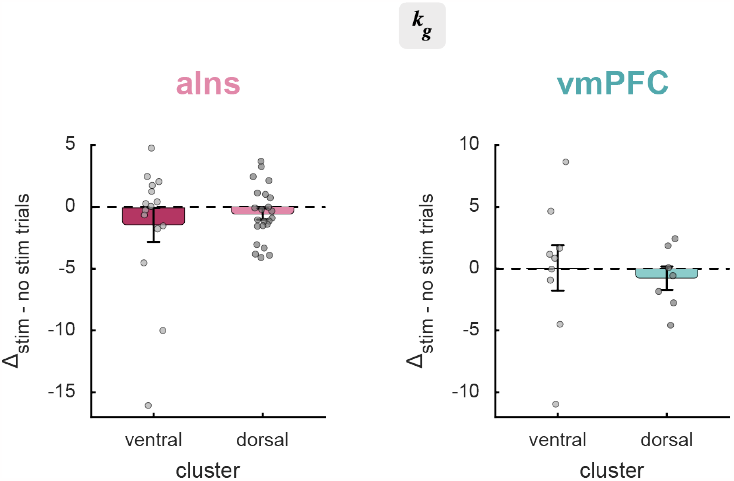
Effects of iES on participants’ sensitivity to potential gains (k_g_). Difference in model weight (i.e., posterior parameter) k_#_ between fits to trials with and without intracranial electrical stimulation in anterior insula (aIns; left) and ventromedial prefrontal cortex (vmPFC; right) clusters. Dots represent individual data. Bars and error bars respectively represent means and SEM across stimulation sites.

**Supplementary Figure 3:**
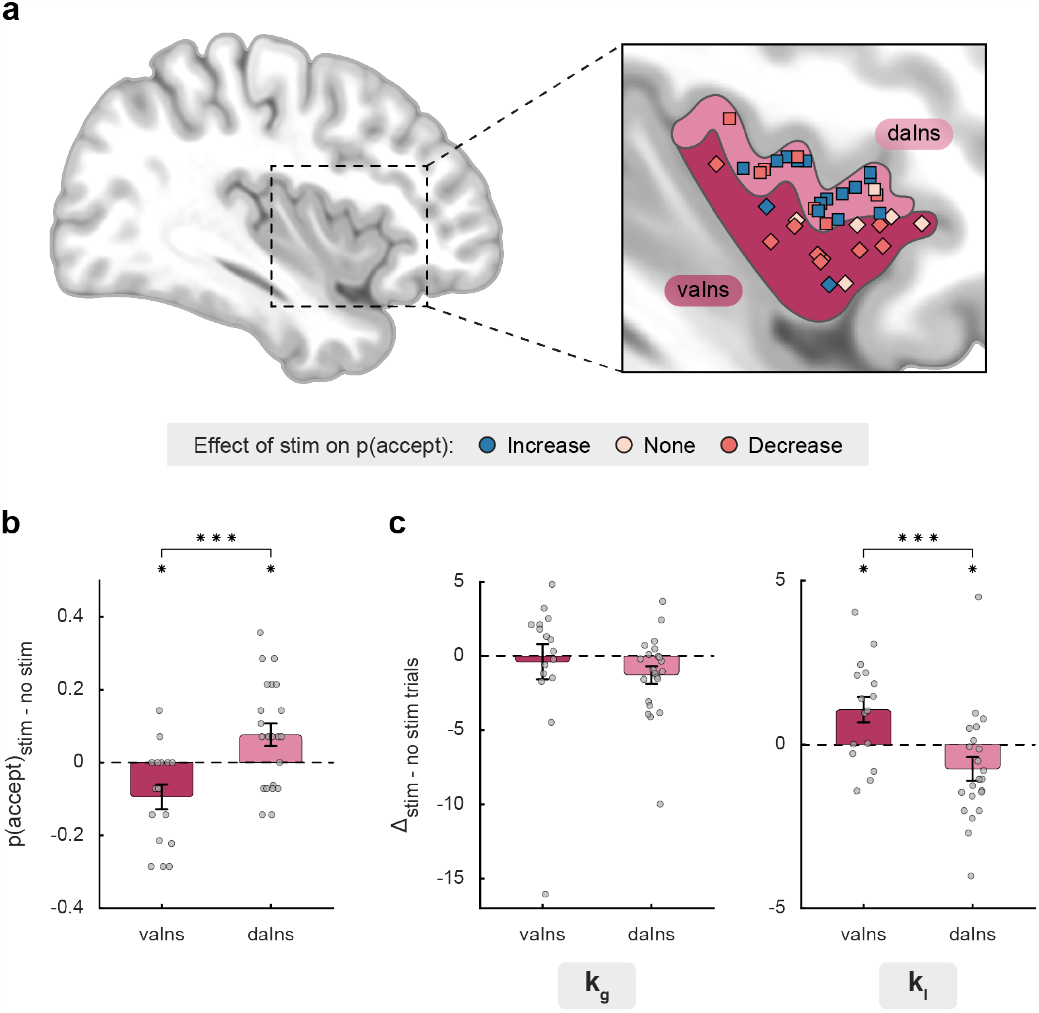
Stimulation effects on acceptance probability in ventral vs. dorsal anterior insula. **(a)** Position of the stimulation sites according to Destrieux’s parcellation scheme. Each electrode was manually positioned on an MNI template according to the individual anatomy of the subjects (see **Supplementary Figure 1** for the location of each site on individual anatomies). The ventral anterior insula (vaIns; diamond) corresponds to the short insular gyri and anterior circular sulcus of the insula in the Destrieux parcellation scheme, whereas the dorsal anterior insula (daIns; square) corresponds to the superior circular sulcus of the insula. Note that this partitioning based on individual anatomy resulted in groupings very close to the previous clusters (over 76% similarity between the two methods). **(b)** Effects of intracranial electrical stimulation (iES) on subregions of the anterior insula. Dots represent individual difference in acceptance probability between trials with and without iES for each stimulation site. **(c)** Computational mechanism of the effect of stimulation of subregions of the anterior insula on decision making. Dots represent the difference in model weights, kg and kl, between fits to trials with and without iES for each stimulation site in the vaIns and daIns. In b and c, bars and error bars respectively represent means and SEM across stimulation sites. *p < 0.05, **p < 0.01, ***p < 0.001.

**Supplementary Figure 4:**
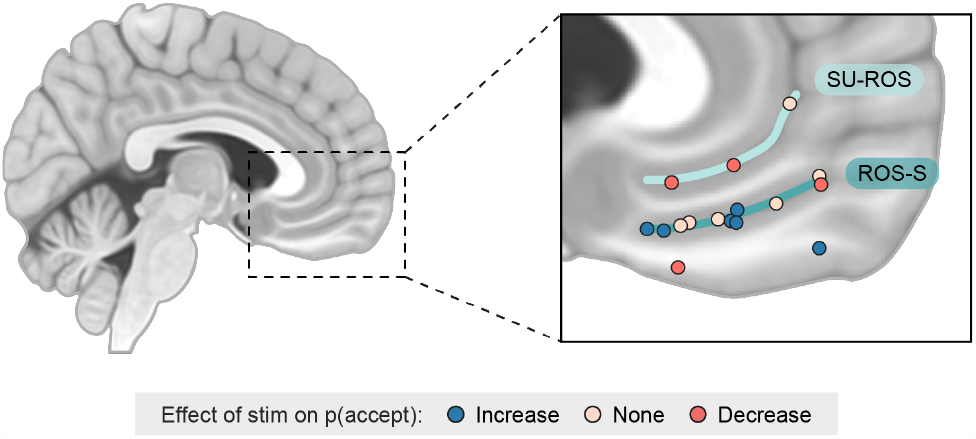
Stimulation effects on acceptance probability as a function of vmPFC anatomy. Position of the stimulation sites according to the two main sulci of the ventromedial prefrontal cortex (vmPFC), the suprarostral sulcus (SU-ROS) and the superior rostral sulcus (ROS-S). Each electrode was manually positioned on an MNI template according to the individual anatomy of the subjects (see **Supplementary Figure 1** for the location of each site on individual anatomies).

**Supplementary Figure 5:**
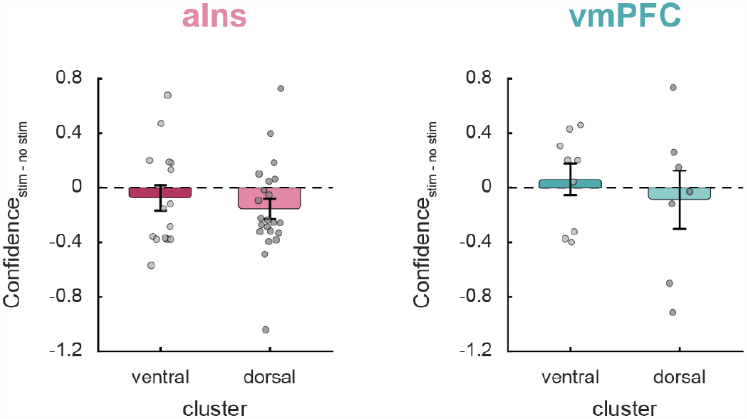
Stimulation effects on confidence (at clusters level). Dots represent individual difference in confidence ratings between trials with and without intracranial electrical stimulation for each stimulation site in the anterior insula (aIns; left) and ventromedial prefrontal cortex (vmPFC; right) clusters. Bars and error bars respectively represent means and SEM across stimulation sites.

**Table S1:**
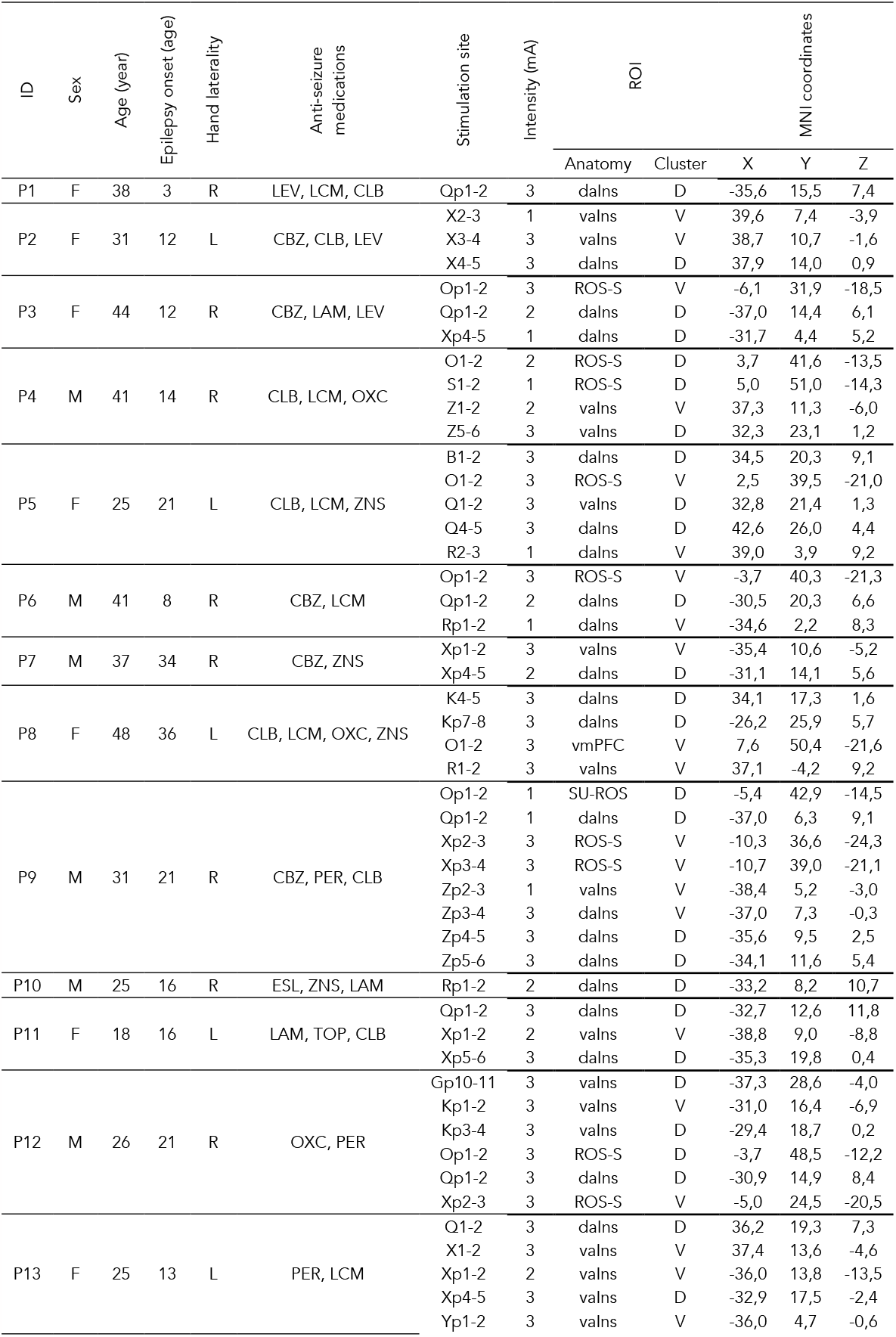
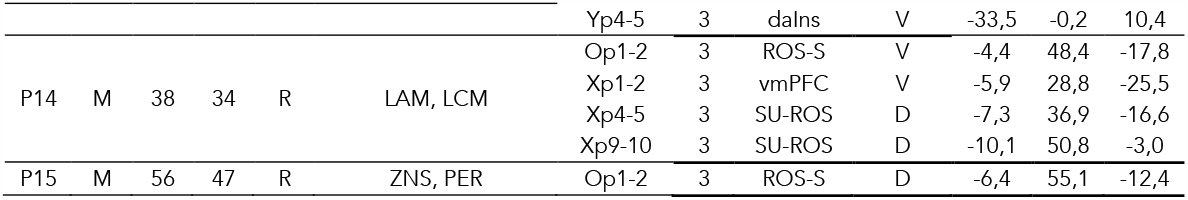
Demographical data. CBZ: carbamazepine; CLB: clobazam; D: dorsal; daIns: dorsal anterior insula; F: female; L: left; LCM: lacosamide; LEV: levetiracetam; LOR: lorazepam; M: male; OXC: oxcarbazepine; PER: perampanel; R: right; ROS-S: superior rostral sulcus; SU-ROS: suprarostral sulcus; V: ventral; vaIns: ventral anterior insula; vmPFC: ventromedial prefrontal cortex; ZNS: zonisamide.

## Material & Methods

### Patient selection

Subjects were fifteen patients suffering from pharmaco-resistant focal epilepsy and candidates to surgical treatment (34.9 ± 2.7 years old, 7 females, 5 left-handed, see demographic details in **Table S1**). These patients underwent a monitoring of their neural activity at the Epilepsy Monitoring Unit of Grenoble University Hospital (Grenoble, France) using stereotaxic implanted multilead deep electrodes (sEEG) as part of their pre-surgical evaluation meant to localize the epileptogenic zone that could not be identified through noninvasive methods. Electrode implantation was performed according to routine clinical procedures with all targeted structures selected strictly according to clinical considerations for the pre-surgical evaluation with no reference to the current study. Patients were included in the study if they had electrodes implanted in at least one brain regions of interest (i.e., aIns and/or vmPFC), and if they were willing and able to perform the choice task. All patients were taking anti-seizure medications (see Supplementary Table S1), some of which were reduced or stopped before stimulation sessions on clinical grounds. Exclusion criteria were age under 18 and complete inability to speak French (to prevent improper execution of the behavioral task due to misunderstanding of task instructions). All patients gave oral informed consent before their inclusion in the present study and signed a non-opposition file in the context of the MAPCOG sEEG study (IdRCB: 2017A03248-45), approved by the Ethics Committee.

### Electrodes implantation and location

Depth electrodes were implanted using robot-assisted sEEG electrode implantation technique (ROSA robot). Fifteen to eighteen semi-rigid electrodes were implanted per patient. Each electrode had a diameter of 0.8 mm and, depending on the target structure, contained 10-18 contact leads of 2 mm wide and 1.5 mm apart (Dixi Medical, Besançon, France). Stimulation sites were chosen in consultation with neurologists for their location in one of our two regions of interest (aIns or vmPFC). To confirm their exact location, the electrodes were also anatomically labeled by co-registering a pre-operative anatomical magnetic resonance imaging (MRI, 3D T1 contrast) with a post-operative computed tomography (CT) scan obtained for each patient, using the IntrAnat Electrodes software (Deman et al., 2018). Subregions of the aIns were identified using the Destrieux atlas (Destrieux et al., 2010). The ventral anterior insula (vaIns) was defined as the short insular gyri (parcel 18; G_insular_short) and anterior circular sulcus of the insula (parcel 47; S_circular_insula_ant), whereas the dorsal anterior insula (daIns) was defined as the superior circular sulcus of the insula (parcel 49; S_circular_insula_sup). Note that the superior circular sulcus of the insula also encompasses part of the posterior insula, but because our stimulation sites were selected anteriorly to be in the anterior insula (i.e., anterior to the central sulcus), all sites were in the anterior part of this parcel (y > -0.2 in MNI space). The vmPFC stimulation sites matched the corresponding parcel of the MarsAtlas parcellation scheme (Auzias et al., 2016). Identification of vmPFC sulci on individual anatomies was done manually based on the literature standard (Lopez-Persem et al., 2019).

### Intracranial electrical stimulation

Intracranial electrical stimulations (iES) were applied between two contiguous contacts located in a region of interest. Bipolar stimuli were delivered on a pair of contacts (defined as a stimulation site) using a constant current rectangular pulse generator designed for a safe diagnostic stimulation of the human brain (Micromed, Treviso, Italy), according to parameters used in clinical procedures and proven to produce no structural damage. High-frequency stimulation at 50 Hz, with a pulse width of 0.5 ms and an intensity of 1, 2, or 3 mA, was applied in a bipolar fashion during a 5 s period on a stimulation site. For a given stimulation site, the stimulation intensity was determined as the highest intensity devoid of clinical symptoms during previous clinical stimulation sessions as systematically done in our routine procedure.

### Stimulation sessions

Each session of the experiment corresponded to the stimulation of a specified stimulation site at a given intensity. Thus, the number of sessions performed by each patient was determined by the number of contact pairs available in regions of interest. Each experimental session consisted of 28 trials of a behavioral task alternating between stimulation (n = 14) and non-stimulation (n = 14) trials (Fig. 1b). The first trial of the session was randomly assigned to either stimulation or non-stimulation to maintain patients’ blindness on experimental conditions. Stimulations were triggered manually and iEEG activity was monitored in real-time to detect stimulation-induced after discharges and electrographic seizures, and to stop the session immediately if necessary. The between trials time interval was also kept above 30 seconds, so that two iES remained separated by a minimum interval of one minute. In stimulation trials, the stimulation was initiated about 1 s before the trial onset.

### Behavioral task

Presentation of visual stimuli and acquisition of behavioral data were performed on a PC using custom Matlab scripts implementing the PsychToolBox libraries (Brainard, 1997). All subject responses were done with a gamepad (Logitech F310S) using both hands. Before the experiment, patients were informed that various brain regions would be stimulated, but they were blind to experimental conditions, including stimulation parameters, the brain region being tested, and whether stimulation was active. Each trial consisted of a choice task combined with confidence rating and a challenge (Fig. 1).

#### Choice task

The choice task began with the presentation of an offer consisting of three attributes: a gain prospect (represented by a bunch of 10-cent coins, range: 1-5*€*), a loss prospect (represented by crossed out 10-cent coins, range: 1-5*€*) and the upcoming challenge difficulty (represented by a target window corresponding to approximately 50% theoretical success). Challenge difficulty was displayed on the center of the screen (see training section for further details about how difficulty was adjusted to each subject). Subjects were asked to accept or reject this offer by pressing a left or right button depending on where the choice option (“yes” or “no”) was displayed. Subjects’ choice determined the amount of money at stake: accepting meant that they would eventually win the gain prospect or lose the loss prospect based on their performance in the upcoming challenge, whereas declining the offer meant playing the challenge for minimal stakes (winning 10 cents or losing 10 cents). The sequence of trials was pseudo-randomized such that all delta values, computed as the difference between gain and loss prospects, continuously sampled along ten intervals ([-40 -30], ]-30 -20], ]-20 -10], ]-10 -5], ]-5 0], ]0 5], ]5 10], ]10 20], ]20 30],]30 40]), were displayed for one patient during a session, with the four medium intervals (]-10 -5], ]- 5 0], ]0 5] and ]5 10]) presented twice (n = 8 out of 14 trials) to maximize the occurrence of difficult choices. The positions of gain and loss prospects were randomly determined to be either displayed on top or bottom of the screen and similarly, the choice options (“yes” or “no”) were randomly displayed on the left or right. Stimulation and non-stimulation trials were strictly identical in one session, such that subjects served as their own control. Patients had a free time delay to accept or decline the offer. If they declined the offer, a 250 ms screen displayed the new offer (a minimal stake of 10 cents). Thus, the challenge was performed regardless of the choice answer to prevent patients from eventually rejecting more offers to decrease experiment duration.

#### Confidence ratings

Before performing the challenge, patients were asked to rate their confidence in winning the challenge by answering the following question “Do you think you will win?”. Patients had a free time delay to answer by moving a cursor from left (not sure) to right (sure) along a continuous visual analog scale (100 steps) with left- and right-hand response buttons. The initial position of the cursor on the scale was randomized to avoid confounding confidence level with movements’ quantity.

#### Challenge

The challenge started right after confidence confirmation: a ball appeared on the left of the screen and moved, horizontally and at constant speed, towards screen center. Patients were asked to press the confirmation button when they thought the ball was inside the box displayed at screen center (i.e., the target window). The ball always reached the center of the target after 1s. Thus, the size of the target window represented the margin of error tolerated in patient reaction time (target: 1s after the movement onset of the ball). Unbeknownst to the patients, the success rate was maintained at about 50% by decreasing (if the challenge was successful) or increasing (if the challenge was unsuccessful) the tolerated margin of error by 1% theoretical success after each trial. Importantly, the moving ball could only be seen during the first 500ms (half of the trajectory), and patients had to extrapolate the last 500ms portion of the ball’s trajectory to assess whether the ball was inside the target. Finally, feedback of 1 second was given to the patients about their payoff after the challenge. Also note that, to improve patients’ motivation to perform the task as accurately as possible, the total amount of money earned by the patients during a session (calculated by adding gains and losses across all trials) was displayed at the end of a session.

#### Training

Before the main experiment, a training - divided into three steps - familiarized subjects with all sub-parts of the task. In the first step, they were familiarized with the challenge by performing 30 to 80 trials of it. Unbeknownst to the subjects, the size of the bar was decreased after each success and increased after each error by one pixel, so that their performance statistically converged at 50% success. This first task was completed when subjects’ performance stabilized (most often before less than 80 trials). Each training trial was followed by feedback informing whether the challenge was successful (“ok” in green) or missed (“too slow” or “too fast” in red). In the second step, subjects completed 64 trials of the full choice (i.e., the challenge was always preceded by an offer), and feedback on the money won/lost in the trial was displayed at the end of each trial. The goal was to train patients to properly integrate the dimensions of the offer (gains and losses) when making their choice. Finally, the third and last part of the training (8 trials) was completely similar to the main task to allow subjects to familiarize with confidence ratings.

Another purpose of the training was to tailor the difficulty of the challenge to each subject’s abilities. To do this, we calculated the mean and standard deviation of challenge performance across all training trials (including all three steps) for each subject, assuming that errors were normally distributed. We then computed an individual tolerated margin of error, corresponding to a theoretical success of 55%, to be used on the first trial of a session of the main task. Among patients, the tolerated margins of error corresponding to 55% theoretical success ranged from [± 40 ms] in the most precise patient to [± 206 ms] in the less precise one.

### Behavioral analysis

Statistical analyses were performed with Matlab Statistical Toolbox (Matlab R2018b, The MathWorks, Inc., USA), and generalized linear mixed-effect (glme) models were estimated using the “fitglme” function with default parameters that maximize the maximum pseudo-likelihood of the observed data under the model. Post-hoc tests were performed using the Matlab function “coefTest”.

#### Choice behavior

Analysis of choice behavior was performed on non-stimulated trials across all sessions (i.e., stimulation sites) using glme models that included choice as the dependent variable (modeled using a binomial response function distribution, i.e., logistic regression) and (1) challenge difficulty, gain and loss magnitude, or (2) confidence as predictor variables. The models also comprised a full random-effects structure at the subject and stimulation site levels (i.e., intercepts and slopes for all predictor variables). For example, in Wilkinson-Rogers notation, the glme model of the effect of offer dimensions on choices can be written as follows: *choice* ∼ 1 + *gain* + *loss* + *diff* + (1 + *gain* + *loss* + *diff*|*subject*) + (1 + *gain* + *loss* + *diff*|*stimulation site*). For each fixed effect, the estimated coefficient value (β) ± its standard error is reported, as well as the t-statistics (testing that the coefficient is equal to 0) along with its p-value.

#### Spatial gradient of the effect of stimulation on acceptance probability

This glme model included the effect of stimulation on acceptance probability as the dependent variable (coded as -1, 0, or 1), the MNI coordinates x, y, and z as predictor variables, and a full random-effects structure at the subject level (i.e., intercepts and slopes for all predictor variables). For each fixed effect, the estimated coefficient value (β) ± its standard error is reported, as well as the t-statistics (testing that the coefficient is equal to 0) along with its p-value.

#### Interaction effects of stimulation and clusters/subregions on choice/confidence

For each region of interest (ROI), these glme models included (1) acceptance probability across trials of the same stimulation condition, or (2) confidence ratings as the dependent variable. The models also comprised main effects and interaction of the stimulation condition and clusters (or subregions) as predictor variables, a full random-effects structure at the subject level (i.e., intercepts and slopes for all predictor variables), and the main effect of stimulation condition at the stimulation sites level. For example, in Wilkinson-Rogers notation, the glme model of interaction effects between stimulation conditions and clusters on choice can be written as follows: *p*_*accept*_ ∼ 1 + *stim condition* ∗ *cluster ID* + (1 + *stim condition* ∗ *cluster ID*|*subject*) + (1 + *stim condition*|*stimulation site*). The coding of the dummy variables was specified as “effects”, so that the sum of the dummy variable coefficients is equal to 0 (Singmann & Kellen, 2019). The F-value of the interactions, main effects, and post-hoc tests are reported along with the degrees of freedom and corresponding p-value. Please note that one participant did not perform confidence ratings, which resulted in the use of one less stimulation site in the anterior insula to investigate the effects of stimulation on confidence (37 sessions instead of 38).

Because spatial sampling from the anatomically-defined dorsal part of the vmPFC (i.e., the suprarostral sulcus; SU-ROS) was not sufficient, we also tested the effect of iES only on the ventral part of the vmPFC (i.e., the superior rostral sulcus; ROS-S) by removing the subregion predictor from the model, yielding for example: *p*_*accept*_ ∼ 1 + *stim condition* + (1 + *stim condition*|*subject*) + (1 + *stim condition*|*stimulation site*).

### Computational analysis

#### Choice model

Choices were fitted using a published computational framework (Cecchi et al., 2022; Vinckier et al., 2018) on additional behavioral data acquired separately from 10 of 15 subjects (128 or 192 trials of the task without stimulation). Acceptance probability was calculated as a sigmoid function (softmax) of expected utility:

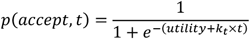

where *k*_*t*_ is a free parameter that accounts for a linear drift with time (trial index *t*) to capture fatigue effects. The utility function is based on expected utility theory where potential gains and losses are multiplied by probability of success (*p*_*s*_) vs. failure (1 – *p*_*s*_):

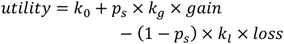

However, distinct weights were used for the gain and loss components (*k*_*g*_ and *kl* respectively), and a constant *k*_0_ was added to capture a possible bias.

The subjective probability of success (*p*_*s*_) was inferred from the target size. The model assumes that subjects have a representation of their precision following a Gaussian assumption, meaning that the subjective distribution of their performance could be defined by its mean (the required 1 second to reach target center) and its width (i.e., standard deviation) captured by a free parameter *σ*. Thus, the probability of success was the integral of this Gaussian bounded by the target window:

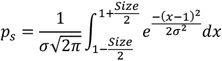

This model was inverted for subjects separately using the Matlab VBA toolbox (available at https://mbb-team.github.io/VBA-toolbox/), which implements Variational Bayesian analysis under the Laplace approximation (Daunizeau et al., 2014).

#### Computational analysis of stimulation effect on choice

The choice model was then run separately with data from trials with and without intracranial electrical stimulation for our 15 subjects. The gain (*k*_*g*_) and loss (*kl*) weights of the expected utility were free to fluctuate, while all other parameters were fixed with posterior means computed with values from the previously inverted model (from the ten subjects who performed an additional behavioral task).

Interaction effects of stimulation condition and clusters/subregions on posterior parameters were tested with the same glme models as the interaction effects of stimulation and clusters/subregions on choice/confidence, but instead of choice/confidence, the dependent variable was the posterior parameters *k*_*g*_ or *kl* obtained with the two trial subsets (with versus without stimulation).

